# Photothrombosis induced cortical stroke produces electrographic epileptic biomarkers in mice

**DOI:** 10.1101/2024.03.01.582958

**Authors:** Dana C. Shaw, Krishnakanth Kondabolu, Katherine G. Walsh, Wen Shi, Enrico Rillosi, Maxine Hsiung, Uri T. Eden, Robert M. Richardson, Mark A. Kramer, Catherine J. Chu, Xue Han

## Abstract

**Objective:** Interictal epileptiform spikes, high-frequency ripple oscillations, and their co-occurrence (spike ripples) in human scalp or intracranial voltage recordings are well-established epileptic biomarkers. While clinically significant, the neural mechanisms generating these electrographic biomarkers remain unclear. To reduce this knowledge gap, we introduce a novel photothrombotic stroke model in mice that reproduces focal interictal electrographic biomarkers observed in human epilepsy.

**Methods:** We induced a stroke in the motor cortex of C57BL/6 mice unilaterally (N=7) using a photothrombotic procedure previously established in rats. We then implanted intracranial electrodes (2 ipsilateral and 2 contralateral) and obtained intermittent local field potential (LFP) recordings over several weeks in awake, behaving mice. We evaluated the LFP for focal slowing and epileptic biomarkers - spikes, ripples, and spike ripples - using both automated and semi-automated procedures.

**Results:** Delta power (1-4 Hz) was higher in the stroke hemisphere than the non-stroke hemisphere in all mice (*p*<0.001). Automated detection procedures indicated that compared to the non-stroke hemisphere, the stroke hemisphere had an increased spike ripple (*p*=0.006) and spike rates (*p*=0.039), but no change in ripple rate (*p*=0.98). Expert validation confirmed the observation of elevated spike ripple rates (*p*=0.008) and a trend of elevated spike rate (*p*=0.055) in the stroke hemisphere. Interestingly, the validated ripple rate in the stroke hemisphere was higher than the non-stroke hemisphere (*p*=0.031), highlighting the difficulty of automatically detecting ripples. Finally, using optimal performance thresholds, automatically detected spike ripples classified the stroke hemisphere with the best accuracy (sensitivity 0.94, specificity 0.94).

**Significance:** Cortical photothrombosis-induced stroke in commonly used C57BL/6 mice produces electrographic biomarkers as observed in human epilepsy. This model represents a new translational cortical epilepsy model with a defined irritative zone, which can be broadly applied in transgenic mice for cell type specific analysis of the cellular and circuit mechanisms of pathologic interictal activity.

**Key Points:**

- Cortical photothrombosis in mice produces stroke with characteristic intermittent focal delta slowing.
- Cortical photothrombosis stroke in mice produces the epileptic biomarkers spikes, ripples, and spike ripples.
- All biomarkers share morphological features with the corresponding human correlate.
- Spike ripples better lateralize to the lesional cortex than spikes or ripples.
- This cortical model can be applied in transgenic mice for mechanistic studies.

## 1. Introduction

Interictal biomarkers in the EEG are routinely used in clinical care to predict seizure risk, localize epileptogenic tissue, and provide rapid assessment of intervention efficacy ^1–4^. Among leading interictal biomarkers, epileptiform spikes, 50 – 200 ms high amplitude voltage fluctuations, are highly specific for epilepsy, but spatially diffuse ^3^. Ripples, brief bursts of 80-250 Hz high-frequency oscillations, are more spatially constrained, but not specific to epilepsy since they are also present during normal cognitive processes ^5,6^. Recently, the combination of these events, spike ripples, has emerged as a biomarker with improved specificity for seizure risk and epileptic tissue ^7,8^. Despite the critical role that interictal epileptic biomarkers play in clinical care, and ongoing efforts to link these events to epileptogenic pathology and mechanisms, little is known about the pathophysiological mechanisms that generate these electrographic biomarkers.

Studies of the cellular and circuit mechanisms of focal cortical epileptic biomarkers require animal models that have defined epileptogenic zones, and ideally transgenic models that permit cell type specific analysis. Of the various animal models of epilepsy, mice that are predisposed to epilepsy due to genetic mutations lack a defined epileptogenic zone required for localizing biomarker studies ^9^. Although systemic or intracranial kainic acid injection models are well established temporal lobe epilepsy models, they do not replicate focal cortical epilepsy ^10^. Similarly, we recently demonstrated that intracranial drug infusion of carbachol or 6-hydroxydopamine in the striatum resulted in pathological spike ripples that correlated with seizure risk in mice ^11^, however this model is limited to the basal ganglia. Electrical kindling via focal administration of electrical stimulation in the brain ^12^ or cobalt wire implantation in the cortex ^13^ can generate symptomatic seizures arising from a defined seizure onset zone, but these models fail to generate spontaneous seizures, suggesting a symptomatic pathophysiology distinct from human cortical epilepsy.

In humans, stroke increases seizure risk and is the most common cause for seizures in elderly populations ^14^. Approximately 11% of stroke patients develop epilepsy within 5 years and the strongest predictor of post-stroke epilepsy is the presence of epileptiform EEG activity ^14–16^. A previous study in rats showed that photothrombosis-induced cortical strokes resulted in epilepsy in 60% of animals ^17^. Here, we examined whether applying photothrombosis in C57BL/6 mice could generate cortical epilepsy biomarkers that are morphologically consistent with human epilepsy biomarkers and localized to injured cortex. The C57BL/6 strain is broadly used for generating the vast number of Cre-transgenic mouse driver lines with Cre-recombinase specifically expressed in defined cell types. Thus, a cortical focal epilepsy C57BL/6 murine model would enable cell-type specific analysis of the cellular and circuit mechanisms of these pathological interictal events.

## 2. Methods

### 2.1. Photothrombosis procedure

All animal procedures were approved by the Boston University Institutional Animal Care and Use Committee. Adult C57BL/6 mice (6 male, 2 female; strain number 000664, Jackson Laboratory) were included in this study. To induce focal stroke, we applied photothrombosis on the cortex similar to that described previously ^17^. Briefly, Rose Bengal (RB, catalog number: 330000, Millipore Sigma) stock solution (8 mg/mL) was freshly prepared by dissolving RB powder in sterile saline (catalog number: NC9054335, Fisher Scientific) on the day of photothrombosis. Mice were first injected subcutaneously with sustained release buprenorphine (3.25 mg/kg, Ethiqa XR, catalog number: 072117, Covetrus) preoperatively. Under isoflurane anesthesia and with the surgical lamp on (KL 200 LED, Leica), we exposed the skull and marked the primary motor cortex (M1) bilaterally (AP: +0.74 mm, ML: ±1.5 mm) to guide later electrode implantation as detailed below. We then injected freshly prepared RB stock solution (40 mg/kg) intraperitoneally (n = 6) or retro-orbitally (n = 2) and waited for 15 minutes to allow for diffusion of RB into the brain with the surgical lamp off to avoid unwanted photothrombosis. We induced photothrombosis anterior to the electrode site, centered at (AP: approximately +1.5 mm, ML: ±1.5 mm). To ensure localized stroke, we covered the surgery lamp with a tin foil mask that had a pin hole of ∼0.5 mm in diameter, and then directed the light onto the exposed skull through the pin hole for 10 minutes. Upon the completion of photothrombosis, the surgical lamp was turned off. With room lights on, a metal head bar was secured on the skull near lambda using metabond (catalog number: 553-3484, Patterson Dental) and dental cement (catalog number: 51459, Stoelting). Upon the completion of surgery, mice were placed in their home cage positioned on a heating pad in a shaded area for at least one hour.

### 2.2. Electrode implantation

Electrode implantation was performed 5-7 days after photothrombosis. Electrodes were made of 0.005” diameter insulated stainless-steel wire (catalog number: 005SW/30S, PlasticsOne) soldered to a 0.016” diameter gold plated dip pin (catalog number: 853-93, Mill-Max Manufacturing Corporation). All mice were implanted with four electrodes (two per hemisphere) to enable within-hemisphere bipolar referencing during data analysis. In each hemisphere, one electrode targeted M1 (AP: +0.74 mm, ML: ±1.5 mm, depth: -1.1 mm) and a second electrode was placed a few millimeters away in cortex (AP:+1.24 mm, ML: ±1.5 mm, depth: -1.2 mm), striatum (AP: +0.24 mm, ML: ±3.0 mm, depth: -3.3 mm), or thalamus (AP: -1.0 mm, ML: ±1.0 mm, depth: -3.5 mm). See Fig 1A for a schematic of the electrode placements and Supplementary Table 1 for electrode target details for each mouse. Under isoflurane anesthesia and after administration of sustained release buprenorphine (3.25 mg/kg, Ethiqa XR, catalog number: 072117, Covetrus), we performed a small craniotomy at each site and lowered the electrodes slowly to the targeted coordinates, and then secured the electrodes on the skull using Metabond (catalog number: 553-3484, Patterson Dental) and dental cement (catalog number: 51459, Stoelting). Additionally, two stainless-steel flathead screws (F000CE0094, J.I. Morris Company) were placed through the skull over the cerebellum to serve as ground and reference during recordings.

**Table 1.** Summary table for the distributions of spike, ripple, and spike ripple rates (event/min) in both the non-stroke (NS) and stroke (S) hemispheres for each of the seven experimental mice across all sessions. These values are computed from the un-transformed distribution (i.e., not in a log_10_ scale). This table includes median, first quartile (Q1), and third quartile (Q3) descriptive statistics for the distributions as well as the *p*-values of the one-sided Wilcoxon signed rank test between the respective stroke and non-stroke hemispheres for each mouse.

| Mouse ID | NS Side: Median (Q1, Q3) | S Side: Median (Q1, Q3) | Wilcoxon Signed Rank Test |
| --- | --- | --- | --- |
| *Spike* |  |  |  |
| 1 | 0.22 (0.12, 0.40) | 4.92 (4.41, 5.45) | $p = 9.766\text{e-}4$ |
| 2 | 0.10 (0.03, 0.20) | 0.12 (0.07, 0.29) | $p = 0.281$ |
| 3 | 0.11 (0.03, 0.20) | 1.56 (1.47, 2.19) | $p = 0.002$ |
| 4 | 0.75 (0.68, 0.86) | 1.23 (1.02, 1.42) | $p = 0.016$ |
| 5 | 0.68 (0.36, 1.21) | 0.12 (0.07, 0.45) | $p = 1.000$ |
| 6 | 0.28 (0.20, 0.37) | 1.33 (1.09, 1.56) | $p = 0.002$ |
| 7 | 0.33 (0.23, 0.42) | 0.89 (0.82, 1.41) | $p = 2.923\text{e-}4$ |
| ***Ripple*** |  |  |  |
| 1 | 2.71 (1.39, 3.90) | 0.59 (0.53, 0.80) | $p = 0.998$ |
| 2 | 7.35 (4.70, 7.97) | 6.86 (4.87, 7.60) | $p = 0.719$ |
| 3 | 1.24 (1.04, 1.88) | 1.31 (1.22, 1.80) | $p = 0.545$ |
| 4 | 5.53 (4.96, 6.59) | 0.70 (0.65, 0.85) | $p = 1.000$ |
| 5 | 1.16 (0.86, 1.49) | 1.31 (0.88, 1.69) | $p = 0.109$ |
| 6 | 4.07 (3.38, 5.34) | 0.58 (0.55, 0.68) | $p = 1.000$ |
| 7 | 5.75 (5.39, 6.30) | 3.21 (2.61, 4.07) | $p = 0.999$ |
| ***Spike ripple*** |  |  |  |
| 1 | 0.00 (0.00, 0.00) | 2.09 (1.79, 2.34) | $p = 9.766\text{e-}4$ |
| 2 | 0.00 (0.00, 0.00) | 2.31 (1.76, 3.20) | $p = 0.016$ |
| 3 | 0.003 (0.00, 0.03) | 0.45 (0.36, 0.67) | $p = 0.002$ |
| 4 | 0.13 (0.02, 0.28) | 0.82 (0.36, 1.00) | $p = 0.031$ |
| 5 | 0.06 (0.03, 0.09) | 0.61 (0.36, 0.68) | $p = 1.605\text{e-}4$ |
| 6 | 0.07 (0.04, 0.25) | 1.06 (0.85, 1.27) | $p = 1.071\text{e-}4$ |
| 7 | 0.04 (0.01, 0.08) | 0.55 (0.41, 0.86) | $p = 2.412\text{e-}4$ |

**Fig. 1.**
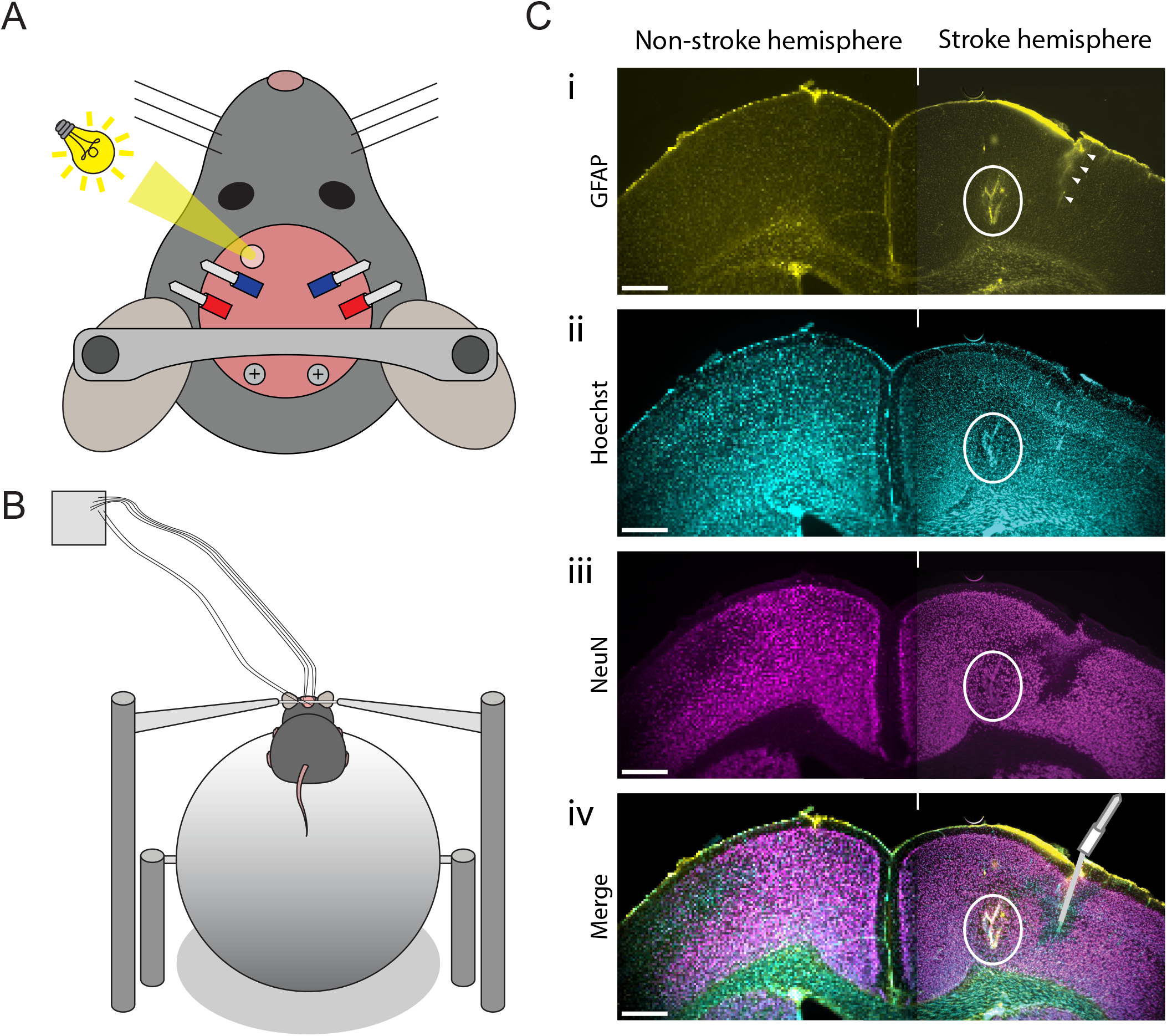
Experimental setup, data collection, and immunohistochemistry. (A) A schematic of the photothrombosis procedure and electrode placement. Light illumination (yellow beam) was directed to a 0.5mm diameter area of the skull (pale pink circle) to generate a stroke in the motor cortex, anterior to the electrode sites. Electrodes (red and blue) were implanted bilaterally, one pair in the stroke hemisphere and the other pair in the non-stroke hemisphere. Animals were also affixed with a head bar and ground and reference screws over cerebellum (gray circles with black plus signs). (B) Schematic of LFP data collection in awake, head fixed mice, freely navigating on a treadmill made of a spherical ball that allows for volitional forward movement. (C) Example of immunofluorescence images from coronal brain slices around the stroke site (marked by white circle), showing GFAP immunofluorescence (i), Hoechst DNA staining (ii), NeuN expression (iii), and a merge of all three (iv). White arrowheads in i indicate gliosis around an electrode tract, with a schematic of electrode placement shown in iv. Scale bars are 0.5 mm.

### 2.3. LFP data collection and analysis

Local field potential (LFP) recordings were collected while mice were head-fixed and freely moving on a treadmill made of a ball pinned along the center axis (Fig. 1B), as described previously ^18^. Wideband LFP was collected at 40 kHz using a Plexon OmniPlex system (Plexon Inc, Dallas, TX). Each mouse was recorded for up to one hour per day, 1-3 times per week over a period of up to 3 months starting at least 15 days post-stroke. To better capture possibly sparse, delayed epileptic biomarkers, we performed 4 extended recording sessions, at least one day apart, with each session lasting for 6 hours for a total of 24 hours at least 3 months post-stroke. See Supplementary Table 1 for detailed information on the number of recording sessions and total durations for each mouse.

LFP power spectra were estimated from 5 second intervals (no overlap, fast Fourier transform with a Hanning taper) and delta power calculated as the summed power across the delta frequency range (1 – 4 Hz). We then calculated the delta power ratio as logarithm base 10 of the ratio of the delta power in the stroke hemisphere over the non-stroke hemisphere, resulting in a distribution of delta power ratios across all recordings for each mouse.

For details on the automated and expert classified biomarkers, please see Supplemental Materials.

### 2.4. Histology

In one mouse (ID: Mouse 1), stroke was confirmed using 2,3,5-triphenyltetrazolium chloride (TTC, catalog number: T8877-25G, Sigma-Aldrich) staining that differentiates viable and necrotic tissue^19^. Briefly, this mouse was perfused intracardially with 0.01M phosphate buffered saline (PBS). The brain was quickly extracted, sectioned into 2 mm slices and placed in 2% TTC in PBS at room temperature for 30 minutes. Immediately after incubation, slices were visually examined and stroke site was confirmed by pale white color of the necrotic tissue, in contrast to the bright red color of viable tissue.

For the other seven mice, we marked electrode locations by passing monophasic electrical current at 100 µA for 60 seconds using a constant current stimulator (Model 2100/4100; A-M Systems) between each electrode and the ground screw to generate electrolytic lesions while mice were anesthetized. Immediately following electrolytic lesions, mice were intracardially perfused with 4% PFA and postfixed for approximately 24 hours. Histology was performed following similar procedures as described previously^20,21^. Briefly, brains were submerged in 30% sucrose solution, and sliced into 50 µm sections using a microtome (Leica SM2010R) and stored in cryoprotectant (1% Polyvinylpyrrolidone, 30% Sucrose, 30% Ethylene Glycol). 7 – 10 brain slices (300 µm apart) spanning the entire M1 area in each mouse were stained with antibodies against NeuN (1:1000, catalog number: ab177487, Abcam) followed by Alexa Fluor 568 goat anti-rabbit (1:1000, catalog number: A11011, Thermo Fisher Scientific), GFAP (1:1000, catalog number: G3893, Sigma Aldrich) followed by Alexa Fluor 488 goat anti-mouse (1:1000, catalog number: A11001, Thermo Fisher Scientific), and Hoechst 33342 (1:1000, catalog number: 62249, Thermo Fisher Scientific). Brain slices were then mounted with ProLong Diamond Antifade Mountant solution (catalog number: P36961, Thermo Fisher Scientific), and imaged using a widefield fluorescence microscope (Leica DM IL LED inverted microscope). Based on the histology validation, one mouse was excluded from our study as the electrode targeting M1 was placed in necrotic tissue. LFP recordings confirmed a lack of physiological signals on this electrode. Thus, final analysis included 7 mice. All stroke sizes were noted to approximate the size of the pin hole (∼0.5 mm in diameter).

### 2.5. Statistical analyses

To compare LFP delta power between the stroke hemisphere and the non-stroke hemisphere across all recordings for each mouse, we performed a nonparametric bootstrapping procedure ^22^. For each mouse, we first computed the proportion of delta ratios greater than zero (delta power in the stroke hemisphere being greater than the non-stroke hemisphere) across all recordings. We then compared the observed proportion to a randomly shuffled distribution. To form the shuffled distribution, we randomly assigned (without replacement) each 5-second interval of summed delta power to the stroke or non-stroke hemisphere and calculated the delta ratio, and then computed the proportion of delta ratios greater than zero during each shuffle. This shuffling procedure was repeated 1000 times to form a distribution of proportions of delta ratios greater than zero, and the *p*-value was calculated based on the amount of proportions computed from the shuffled distributions that were greater than the observed proportion ^22^. We performed this bootstrap procedure separately for each mouse.

To test whether the rates of automatically detected or expert validated spikes, ripples, and spike ripples in the stroke hemisphere were higher than those detected or validated in the non-stroke hemisphere, for each mouse and each biomarker we performed a paired one-sided Wilcoxon signed rank test (Table 1, automatically detected results only). To test whether the rates of validated biomarkers were higher in the stroke hemisphere across all mice, we compared the mean of the differences in biomarker rates between the two hemispheres for each mouse against zero using a one-sided Wilcoxon signed rank test.

To compare the ability of each biomarker (spikes, ripples, and spike ripples) to classify a hemisphere as non-stroke or stroke across mice, we first computed the average rate for each automatically detected or validated biomarker (spike, ripple, and spike ripple) in each hemisphere across all sessions for all mice, resulting in n = 164 automatically detected and n = 14 validated rates for each biomarker. To estimate classification performance, we fit a generalized linear model (binomial distribution with binary outcome “stroke” or “non-stroke” and predictor biomarker rate with a logit link function) to these data, and computed receiver operator characteristic (ROC) curves. The ROC curves and optimal operating points were computed using MATLAB’s *perfcurve* function which were subsequently used to compute sensitivity and specificity.

## 3. Results

### 3.1. Photothrombosis reliably induced a focal stroke in mouse cortex

We induced focal strokes in mouse cortex using a photothrombosis procedure by illumination of an area ∼0.5mm in diameter over the motor cortex (details in Methods). To confirm successful stroke induction, in one mouse, we performed histology at the completion of the study by staining tissue with TTC to visualize the presence of necrotic areas (pale color). In the rest of the mice (n=6), we performed immunohistochemistry for glia marker GFAP and neuronal marker NeuN to detect gliosis and neuronal loss around the stroke sites (Fig. 1C). Across all mice, we found that the photothrombosis procedure reliably induced a localized stroke in the target hemisphere, exhibiting increased GFAP expression and decreased NeuN expression.

### 3.2. Photothrombosis results in characteristic intermittent focal delta (1-4Hz) slowing in the stroke hemisphere

Intermittent or continuous LFP delta oscillations, a.k.a. focal slowing, is frequently observed following ischemic stroke in humans ^23^. Visual inspection of the mouse LFP recordings revealed frequent occurrences of large amplitude LFP delta oscillations in the stroke hemisphere, but not in the non-stroke hemisphere (Fig. 2A). On quantitative analysis, we found increased delta power in the stroke hemisphere compared to the non-stroke hemisphere in each mouse (Fig. 2B, *p* = 0.0005, nonparametric bootstrap for each of the 7 mice). Thus, similar to observations in stroke patients, delta slowing is present in the perilesional cortex ipsilateral to stroke in the mouse photothrombotic model.

**Fig. 2.**
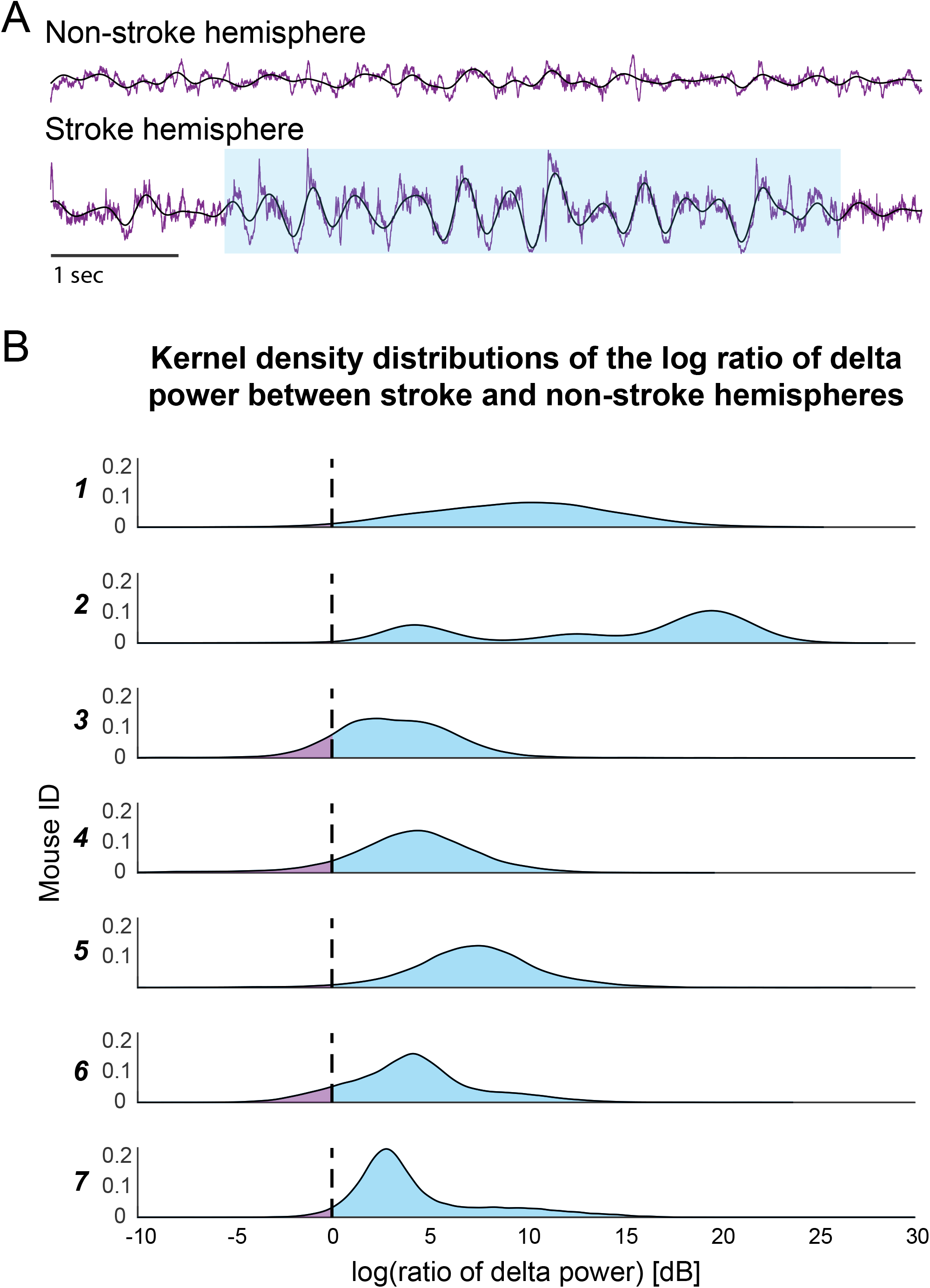
Delta power is increased in the stroke hemisphere due to intermittent focal slowing. (A) An example LFP time series recording (purple trace) demonstrating intermittent focal slowing (blue highlight) in the stroke hemisphere (bottom) and the corresponding LFP in the non-stroke hemisphere (top). Black traces are delta frequency filtered LFPs. (B) Kernel density distributions of the logarithm base 10 of the ratio of delta power in the stroke hemisphere over the non-stroke hemisphere for each mouse. Area under the curve greater than 0 (blue) indicates greater delta power in the stroke hemisphere while area under the curve less than 0 (purple) indicates greater delta power in the non-stroke hemisphere. Delta power was significantly increased in the stroke hemisphere across time in all mice (*p* = 0.0005 for each mouse, nonparametric bootstrap, see Kramer and Eden, 2016).

### 3.3. Automatically detected spikes tend to lateralize to stroke hemisphere

Across recording sessions, we noted many large amplitude events which exhibited similar morphological features as epileptic spikes in humans, typically in the stroke hemisphere, but not always (Fig. 3A). Using an established automatic spike detector used in humans (see Supplementary Materials and Fig. 3Bi), we computed the spike rate for each hemisphere across all mice (Fig. 4A). We found that spike rate in the stroke hemisphere was significantly greater than in the non-stroke hemisphere for 5/7 (71%) of mice (p < 0.05, paired one-sided Wilcoxon signed rank test, see Table 1 for each p-value). Across the population of mice, the mean difference in spike rates between stroke and non-stroke hemispheres was significantly greater than zero (*p* = 0.039, one-sided Wilcoxon signed rank test, Fig. 4D).

**Fig. 3.**
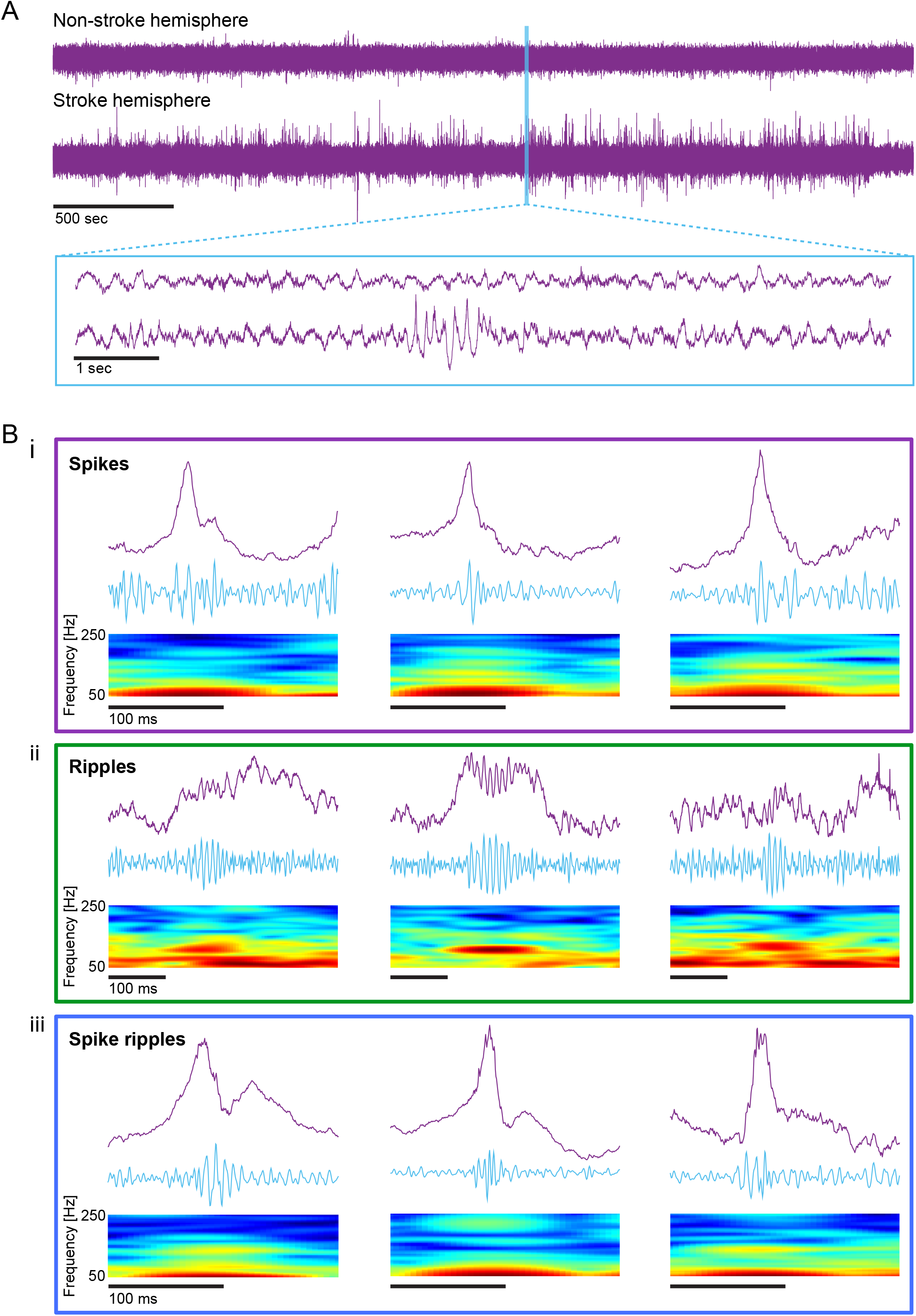
Example interictal events. (A) An example LFP time series recording for a one-hour session from one mouse for both the non-stroke (top purple trace) and stroke (bottom purple trace) hemispheres. The blue box shows a zoom-in of 10 seconds for each hemisphere. (B) Example spikes (i), ripples (ii), and spike ripples (iii). For each example biomarker, the top plot is the raw LFP time series (purple trace), the middle plot is the 100 - 300 Hz bandpass filtered time series (blue trace), and the bottom plot is the spectrogram of the raw time series with frequencies plotted for the 50-250 Hz range.

**Fig. 4.**
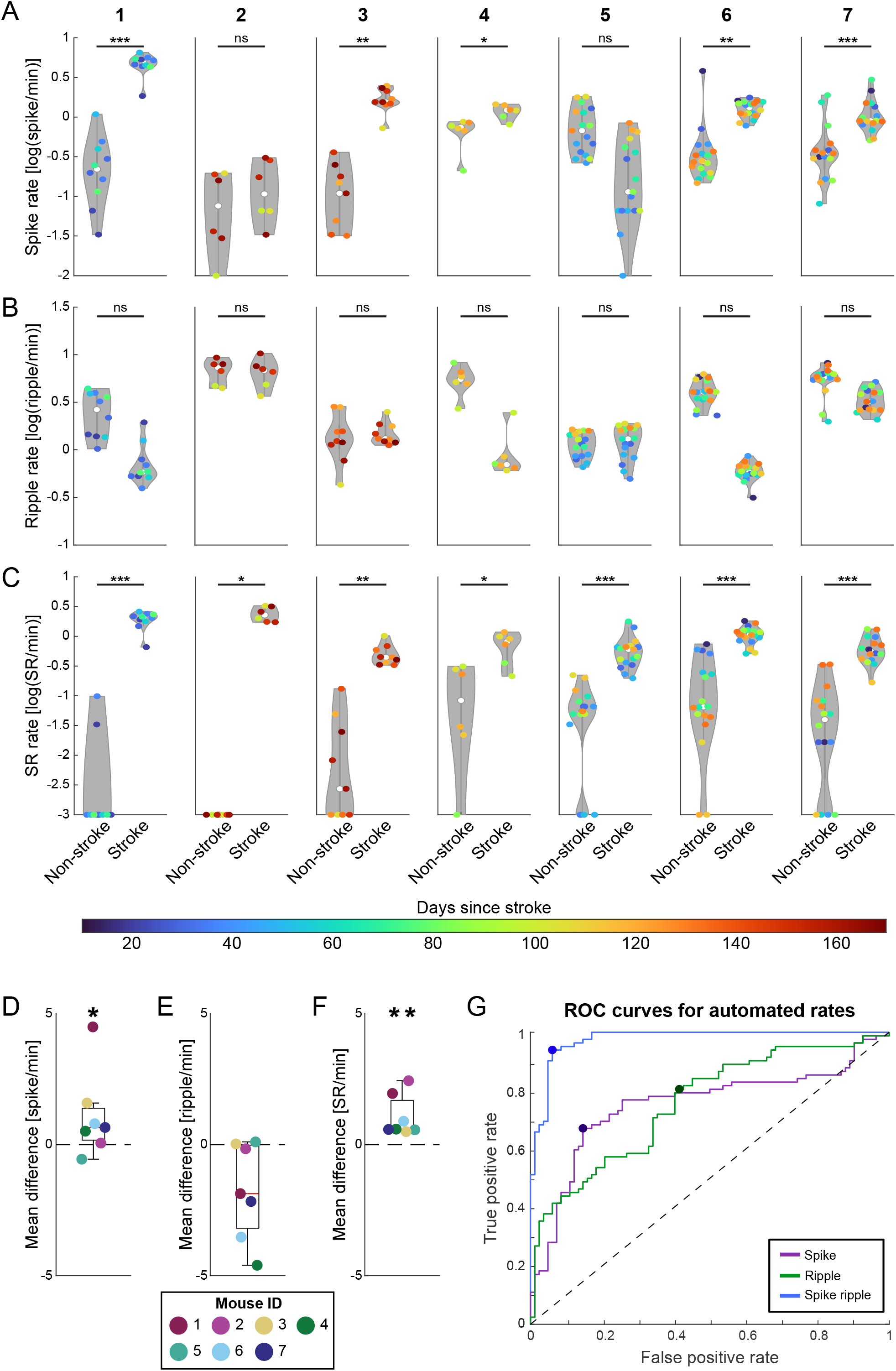
Automatically detected spike ripple rate is consistently increased in the stroke hemisphere compared to spikes or ripples. (A – C) Violin plots for the seven experimental mice (labelled on the top 1-7) of the rates of automatically detected spike (A), ripple (B), and spike ripple (C) biomarkers within the non-stroke and stroke hemispheres. Stroke-induced increase in biomarker rates were determined by paired one-sided Wilcoxon signed rank tests. Each dot represents a recording session, and the color of the dot indicates the number of days since the stroke (ns = not significant, * *p* < 0.05, ** *p* < 0.01, *** *p* < 0.001; see Table 1 for exact *p*-values). For visualization purposes, 0 events/min values were set to -2 in (A) and -3 in (C). (D – F) The mean rate difference between stroke and non-stroke hemispheres calculated across recording sessions in each mouse for spikes (D), ripples (E), and spike ripples (F). The distribution of the mean of the rate differences was compared to zero using a one-sided Wilcoxon signed rank test (* *p* < 0.05, ** *p* < 0.01). Average mean difference in spike, ripple, and spike ripple rates (D: 1.07 + 1.64, E: - 1.75 + 1.85, F: 1.07 + 0.79; average + SD). (G) The ROC analysis curves for all recording sessions across all mice (n = 164 event rates per biomarker) for spike (purple), ripple (green), and spike ripple (blue) rates that were automatically detected. Dashed black line depicts the chance curve. The optimal operating point for each ROC curve is indicated with a filled circle. See Supplementary Table 2 for ROC metrics. SR: spike ripple.

### 3.4. Automatically detected ripples do not lateralize to the stroke hemisphere

On visual inspection, ripples exhibiting similar morphological features as observed in humans were also identified (Fig. 3Bii). Using two different automated approaches to identify ripples (see Supplementary Materials), we found no evidence of increased ripple rate in the stroke hemisphere compared to the non-stroke hemisphere in any of the mice analyzed (*p* > 0.05 in all mice, paired one-sided Wilcoxon signed rank test, see Table 1 for each *p*-value, Fig. 4B and Fig. S1). Across the population of mice, the mean differences in ripple rates between stroke and non-stroke hemispheres were not greater than zero (*p* = 0.98, one-sided Wilcoxon signed rank test, Fig. 4E). Thus, automatically detected ripples do not reliably localize pathologic cortex in this photothrombotic stroke mouse model.

### 3.5. Automatically detected spike ripples lateralize to the stroke hemisphere

Visual review revealed spike ripples that shared consistent morphology with those observed in epileptic patients (Fig. 3Biii). Using a detector validated in human intracranial recordings, automatically detected spike ripple rates were significantly higher in the stroke hemisphere than in the non-stroke hemisphere in each mouse (*p* < 0.05 in each mouse, paired one-sided Wilcoxon signed rank test, Fig. 4C, also see Table 1 for *p*-values). The mean differences in rates between hemispheres across all mice were significantly greater than zero (*p* = 0.008, one-sided Wilcoxon signed rank test, Fig. 4F). Across mice, spike ripples were present and elevated in the stroke hemisphere compared to the non-stroke hemisphere (median difference in spike ripple rate: 0.63 events/min, range 0.26-2.02 events/min) by the first date of post-stroke recording (median 29, range 15-105 days, Supplementary Table 1). Thus, automatically detected spike ripple rate is a reliable, lateralized, early indicator of tissue pathology in this model.

### 3.6. Expert validated ripples and spike ripples lateralize to the stroke hemisphere

To verify the automated detections of spikes, ripples, and spike ripples in mouse LFP recordings, one hour of LFP data from each mouse was reviewed and all detections (855 candidate spikes, 2,346 candidate ripples, and 425 candidate spike ripples) were classified by a human expert blinded to mouse, stroke side, and recording site. In the expert validated events, we found a trend toward higher spike rates in the stroke hemisphere compared to the non-stroke hemisphere (*p* = 0.055, paired one-sided Wilcoxon signed rank test, Fig. 5A). The validated ripple rates were significantly higher in the stroke hemisphere (*p* = 0.031, paired one-sided Wilcoxon signed rank test, Fig. 5B). Finally, the validated spike ripple rates in the stroke hemisphere were higher than the non-stroke hemisphere (*p* = 0.008, paired one-sided Wilcoxon signed rank test Fig. 5C). These results support that each of these interictal biomarkers – spikes, ripples, and spike ripples – are reliably generated by the mouse photothrombotic stroke model ipsilateral to damaged cortex.

**Fig. 5.**
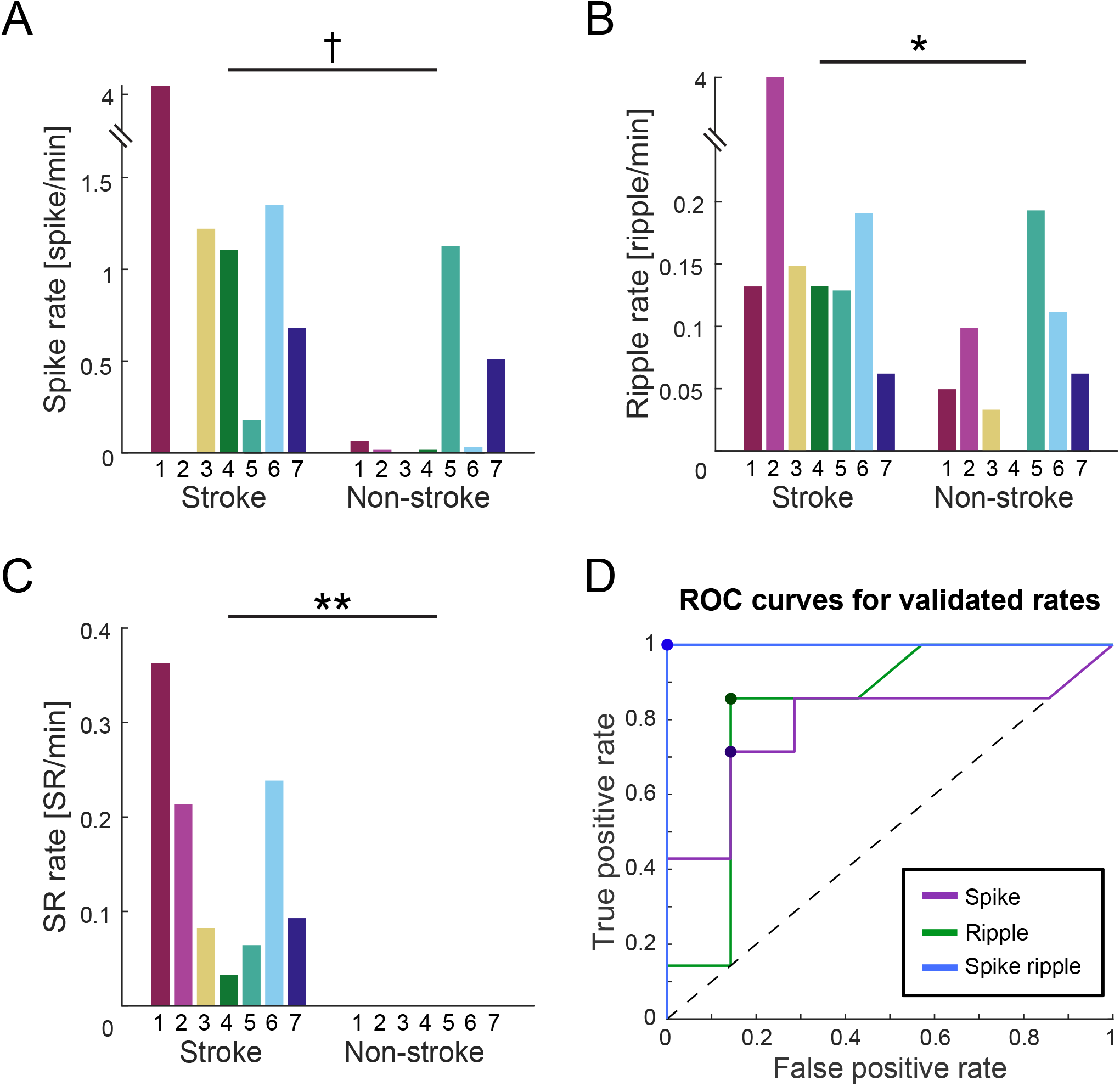
Expert validated spike ripple rate best classifies stroke vs non-stroke hemisphere. (A) The rate of the expert validated spikes during a single one-hour recording session for each of the 7 mice (mouse ID on x-axis). Comparisons between the stroke and non-stroke hemisphere rates were performed using a paired one-sided Wilcoxon signed rank test. (B, C), similar to A, but for expert validated ripples (B) and spike ripples (C) († *p* < 0.10, * *p* < 0.05, ** *p* < 0.01). (D) ROC curves for the expert validated spike (purple), ripple (green), and spike ripple (blue) rates (n = 14 event rates per biomarker). The dashed black line depicts the chance curve and the optimal operating point for each ROC curve is indicated with a filled circle. See Supplementary Table 2 for ROC metrics. SR: spike ripple.

### 3.7. Spike ripple rates better classify the stroke hemisphere than spike or ripple rates

To evaluate the ability for these biomarkers to distinguish the stroke and non-stroke hemispheres, we classified each hemisphere within each session using ROC analysis for automatically detected spikes, ripples, and spike ripples separately (see *Methods*, Fig. 4G, and Supplementary Table 2 for summary metrics). Using the optimal operating point for each automatically detected event type (0.797 spikes/min, 0.314 ripples/min, 0.330 spike ripples/min; Fig. 4G), spike ripples classified the stroke hemisphere from the non-stroke hemisphere with high sensitivity (0.94) and specificity (0.94). In contrast, spikes and ripples yielded lower sensitivities (0.67 and 0.81, respectively) and specificities (0.85 and 0.59, respectively).

ROC analysis using these validated rates again demonstrated that spike ripples were superior to spikes alone or ripples alone at classifying the pathological hemisphere. Using the optimal operating point for each manually validated event type (0.68 spikes/min, 0.13 ripples/min, 0.03 spike ripples/min; Fig. 5D), spike ripples classified the stroke hemisphere with higher sensitivity (1.00) and specificity (1.00) than spikes or ripples (sensitivity 0.71 and 0.86, respectively; specificity 0.86 and 0.86, respectively; Fig. 5D and Supplementary Table 2 for summary metrics).

These results are consistent with recent work in humans demonstrating that automatically detected spike ripples are superior to automatically detected spikes or ripples alone at localizing epileptogenic brain tissue ^8^. Together, these results demonstrate that the photothrombosis-induced stroke mouse model generates robust spike, ripple, and spike ripple biomarkers that share morphological features with epileptic events observed in human EEG and LFP recordings. Further, these results support findings that spike ripples have improved specificity for pathologic tissue compared to spikes or ripples alone.

## 4. Discussion

We demonstrate that photothrombosis-induced cortical focal stroke in mice reliably generates the canonical electrophysiological events observed in human stroke and post-stroke epilepsy, including intermittent focal slowing, spikes, ripples, and spike ripples. Further, we showed that spike, ripple, and spike ripple biomarkers can be detected using automated methods developed for human EEG data, demonstrating this stroke model produces biomarkers with shared morphology observed in human patients. Finally, spike ripples outperformed other biomarkers in reliably lateralizing the stroke and non-stroke hemispheres with high sensitivity and specificity. Thus, this cortical photothrombosis stroke mouse model represents a new system for mechanistic studies of biomarkers associated with cortical epilepsy.

While we observed spikes, ripples, and spike ripples in the damaged neocortex in our mouse model that approximate biomarkers observed in human cortex, spike and ripple events did not lateralize to the damaged cortex when using automated detectors optimized for human recordings. While these results could indicate poor detector performance, spikes maintained limited spatial specificity even after manual classification, and these results are consistent with observations in humans ^3^. Manually classified ripples lateralized to the damaged cortex better than the automated detections, highlighting the challenges in automatically detecting these brief, subtle events. However, hand-marking of ripple events is an especially time-consuming process with poor reliability ^24^. Existing evidence shows that ripples often co-occur with spikes ^7,24–26^, and these combined events are easier to detect than ripples alone using automated approaches. Increasing evidence in humans suggests that these co-occurring events – spike ripples – better predict seizure severity and localize epileptogenic tissue than either spikes alone or ripples alone ^1,7,8,25^. Consistent with these observations in humans, our results indicate that spike ripples reliably indicate pathological tissue in the mouse cortical stroke model.

Approximately 5 – 20 % of stroke patients will develop post-stroke epilepsy, defined by at least two unprovoked seizures after initial injury, with a majority developing epilepsy within the first year after stroke ^14,27,28^. In such patients, the presence of intermittent focal slowing on early EEG monitoring (i.e., within the first 72 hours after stroke) can help indicate the severity of injury ^29–31^ and the presence of interictal spikes can help predict the risk of developing epilepsy ^32,33^. Here, we observed both of these features in our mouse stroke model, where both increased delta oscillations and interictal biomarkers were present in the first recording (as early as 15 days) post-stroke. These data indicate that the photothrombosis model yields robust post-stroke pathology in mice that mimics what can occur in human stroke patients that go on to develop epilepsy.

Our study had several limitations. First, while the results presented here indicate the automated detectors used in humans can detect comparable biomarkers in mice, each detector could be further optimized for improved performance in mice. As the majority of false detections in our dataset were due to artifacts, improved pre-processing or detection strategies that are more robust to movement and lead artifacts could improve detector performance. In addition, given that epileptiform discharges predict epilepsy in humans, here we focused only on interictal biomarkers. Prior work using the photothrombosis model in epilepsy has mainly been performed in rats where electrographic or behavioral seizures have been successfully captured ^17,34,35^. Over the course of this study, we did not capture any electrographic or definite behavioral seizures. Seizures can be rare events and difficult to capture in many mouse models of epilepsy ^36^. Although we performed several monitoring sessions, confirmation of seizures in this focal stroke model would likely require multi-week continuous EEG monitoring. Alternatively, future work could link these biomarkers to seizures by using a mouse strain more prone to seizures ^37,38^. On the other hand, the C57BL/6 strain used in this study supports many transgenic lines that would allow for a more thorough dissection of the cellular networks generating interictal events, such as cell-type specific analysis, than other strains more prone to seizures which do not have such transgenic lines (i.e., CD1 mice).

These results support future work to probe the cellular and network mechanisms underlying spikes, ripples, and spike ripples. Interictal spikes have been proposed to involve both glutamatergic and GABAergic synaptic signaling ^39–42^. Ripples have been proposed to depend on fast-spiking interneurons ^43,44^ or gap-junction coupled pyramidal cells ^45,46^. Using cellular voltage imaging techniques, such as that demonstrated in our recent studies using genetically encoded voltage indicators ^47,48^, future studies could examine the relationship between membrane voltage of specific types of neurons and interictal epileptiform activity in epilepsy. Thus, insight into the pathological mechanisms underlying spike ripples and other biomarkers would be possible with the development of this mouse model that generates these cortical events.

## Conclusions

In conclusion, we developed a mouse model of cortical stroke that yields epileptic biomarkers soon after initial injury that remain chronically prevalent. The co-occurrence of spikes and ripples – spike ripples – are better lateralized to the injured hemisphere than either spikes alone or ripples alone. Moreover, using both automated approaches and expert visual analysis, the spikes, ripples, and spike ripples found in this mouse model share morphological features with those observed in human patients. This model enables future work on interrogating the cellular and circuit mechanisms underlying the emergence of such biomarkers.

## Abbreviations

LFP: local field potential
GFAP: glial fibrillary acidic protein
NeuN: neuronal nuclei

## Data availability

Rodent data used in this analysis is available upon reasonable request from the corresponding authors.

## Author contributions

C.J.C, M.A.K. and X.H. conceived, planned, and supervised the experiments.

K.K and D.S. conducted the experiments. D.S. wrote the first draft of the manuscript. D.S., W.S., K.G.W., E.R., M.H. contributed to data analysis. All authors discussed the results and contributed to the final manuscript.

---

**Fig. S1.**
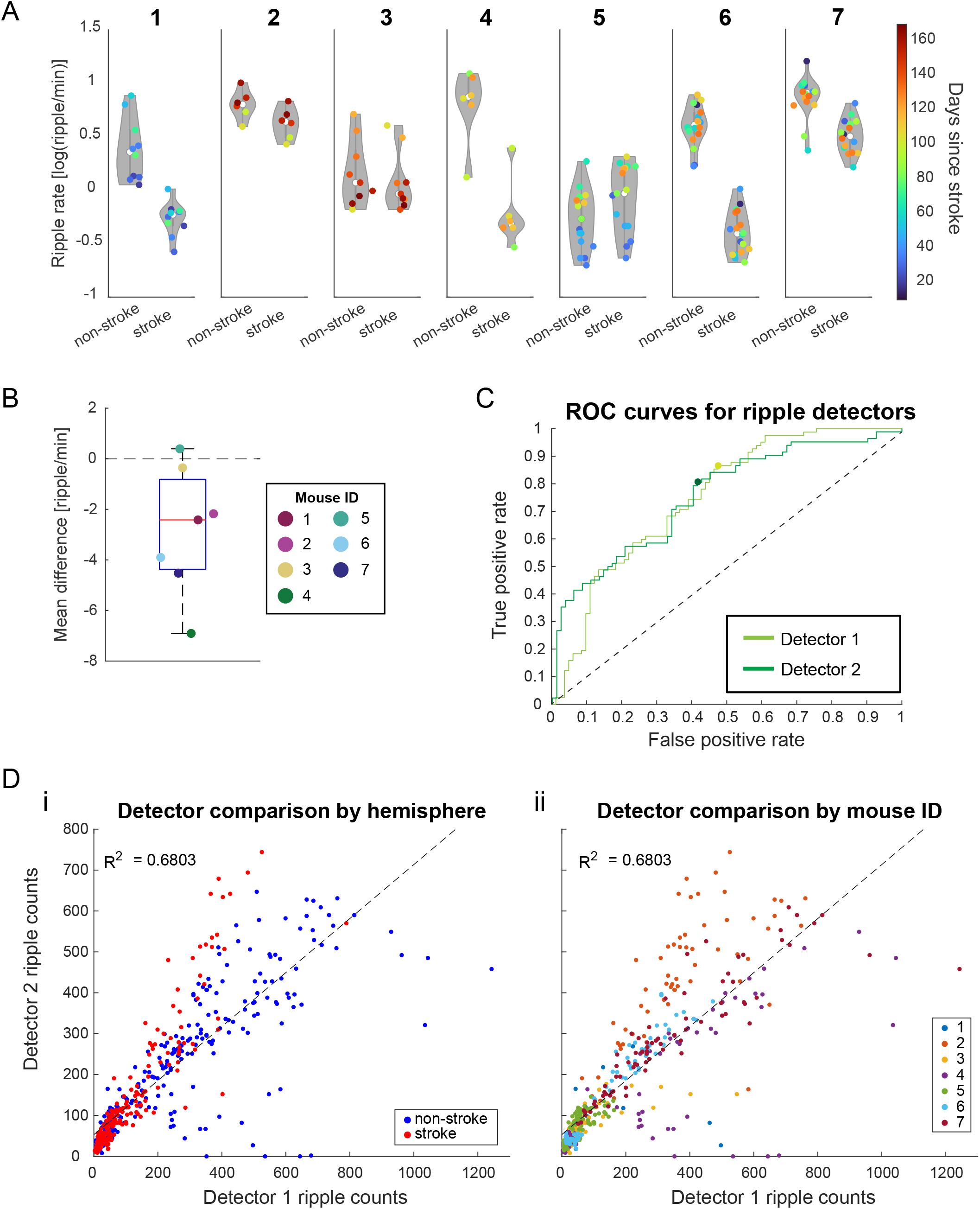
The automatic ripple detector developed by Staba and colleagues yields results consistent with our altered detector from Chu et. al., 2017. (A) Violin plots for ripple rate in each of the seven mice detected by the ripple detector developed by Staba and colleagues (Detector 1) ^49^. The results are consistent with the ripples detected using the altered detector based on Chu et. al., 2017 (Detector 2), i.e., none of the automatically detected ripple rates are increased in the stroke hemisphere relative to the non-stroke hemisphere (p > 0.05 for each mouse, paired one-sided Wilcoxon signed rank test). (B) A boxplot of the mean difference in ripple rates across all sessions. Again, these results are consistent with Detector 2. Average mean difference in ripple rates (−2.84 + 2.51; average + SD, *p* = 0.984, one-sided Wilcoxon signed rank test). (C) ROC curves for Detectors 1 and 2, highlighting their similarity. Optimal operating points are shown as dots on each respective curve. Detector 1: optimal threshold = 0.198 ripples/min, sensitivity = 0.866, specificity = 0.524, AUC = 0.751. Detector 2: optimal threshold = 0.314 ripples/min, sensitivity = 0.805, specificity = 0.585, AUC = 0.756. (D) Comparison of the number of ripples detected by both detectors while visualizing stroke versus non-stroke hemisphere (i) or mouse ID (ii), where each dot is the number of ripples detected in a hemisphere from any of the 82 total sessions (n = 164). A linear regression (dashed black line) between Detector 1 and Detector 2 yielded an adjusted R^2^ = 0.68.

