## Supplementary Methods for "Photothrombosis induced cortical stroke produces electrographic epileptic biomarkers in mice"

**Supplementary Materials**

**Supplementary Methods**

*Detection of spikes, ripples, and spike ripples*

For automated detection of spikes, we utilized the Persyst 14 software Reveal detector ^1^ (Persyst Development Corporation, San Diego, CA, USA). To ensure consistency across mice and hemispheres, the sensitivity of the Reveal detector was set to 50 uV.

For automated detection of ripples, we applied a modified version of the spike ripple detector described in Chu et. al., 2017. Specifically, LFPs were first bandpass filtered between 100 and 300 Hz (equiripple FIR filter of order 168, pass-band 100 to 300 Hz, pass-band ripple 0.1 dB, frequency of the first stop-band of 60 Hz with attenuation 80 dB, and frequency of the second stop-band of 350 Hz with attenuation 40 dB). We then computed the amplitude envelope of the filtered LFP data using the Hilbert transform, and identified time points when the amplitude envelope was in the top 97.5^th^ percentile. If the identified time points were less than 5 ms apart, the corresponding time points were merged. Ripples were then defined as the time points with durations greater than 20 ms, at least 3 zero-crossing, and approximately sinusoidal in shape, defined as having Fano factors computed using the time intervals between zero-crossings being less than 1, as detailed in (Chu et. al., 2017).

As a second measure of ripple detection, we applied the automated detector developed by Staba and colleagues ^2^ as implemented in the MATLAB toolbox RIPPLEAB ^3^, which has been applied previously for the detection of cortical ripples in mice ^4^. Briefly, LFPs were bandpass filtered between 80 and 250 Hz and the root mean square (RMS) amplitude of the filtered data was computed. Within each 600 second interval of LFP data (non-overlapping), RMS amplitudes greater than 3 standard deviations above the mean amplitude of the RMS signal and with a duration of at least 6 ms were automatically detected. Among these detections, only detections with at least 4 peaks (where peaks were defined as exceeding 3 standard deviations above the mean rectified RMS signal) were included ^3^. Any resulting detections separated by less than 10 ms were merged. Results using this second ripple detection method were qualitatively similar to the ripple detector modified from the spike ripple detector reported above (Fig. S1).

For automated detection of spike ripples, we applied a feature-based spike ripple detector previously developed and applied by our group ^5,6^ and available at https://github.com/Mark-Kramer/Spike-Ripple-Detector-Method. Briefly, the detector identifies candidate spike ripple events by detecting a high frequency oscillation (100 – 300 Hz) approximately sinusoidal in shape, with at least three cycles, that co-occur with a large amplitude discharge (Chu et al., 2017).

In addition to automated event detection, a human expert blinded to experimental conditions and mouse ID visually inspected and classified the detected events in a subset of data. In each mouse, the human expert examined all automatically detected spikes, ripples (detected from the altered Chu et. al., 2017 method described above), and spike ripples separately in a single one-hour recording session.

**Supplementary Table 1**

| Mouse ID | Sex | Rose Bengal Injection Site | Electrode Targets (per hemisphere) | First/Last recording session (DPS) | No. recording Sessions | Total LFP recorded (min) |
| --- | --- | --- | --- | --- | --- | --- |
| 1 | M | IP | M1, Thalamus | 20 / 77 | 10 | 390 |
| 2 | M | RO | M1, anterior M1 | 99 / 167 | 6 | 1390 |
| 3 | M | IP | M1, anterior M1 | 105 / 171 | 9 | 1730 |
| 4 | M | IP | M1, Striatum | 84 / 126 | 6 | 1580 |
| 5 | M | IP | M1, Striatum | 29 / 127 | 17 | 1970 |
| 6 | F | RO | M1, anterior M1 | 15 / 133 | 18 | 2180 |
| 7 | F | IP | M1, anterior M1 | 15 / 134 | 16 | 2130 |

**Supplementary Table 1.** Summary table. IP: intraperitoneal; RO: retro-orbital; M1: primary motor cortex; DPS: days post-stroke

**Supplementary Table 2**

| Biomarker | Threshold (events/min) | Sensitivity | Specificity | AUC |
| --- | --- | --- | --- | --- |
| *Automatically detected rates* | | | | |
| Spike | 0.80 | 0.67 | 0.85 | 0.75 |
| Ripple | 0.31 | 0.81 | 0.59 | 0.76 |
| Spike ripple | 0.33 | 0.94 | 0.94 | 0.98 |
| *Validated rates* | | | | |
| Spike | 0.68 | 0.71 | 0.86 | 0.79 |
| Ripple | 0.13 | 0.86 | 0.86 | 0.83 |
| Spike ripple | 0.03 | 1.00 | 1.00 | 1.00 |

**Supplementary Table 2.** Summary metrics for ROC analysis for automatically detected and validated spike, ripple (using the altered Chu et. al., 2017 method), and spike ripple rates. The reported thresholds are the optimal thresholds in events/min and the reported sensitivities and specificities correspond to these optimal thresholds. Area under the curve (AUC) is reported for each biomarker’s ROC curve.
